## Supplementary Figures for "Cellular events of acute, resolving or progressive COVID-19 in SARS-CoV-2 infected non-human primates"

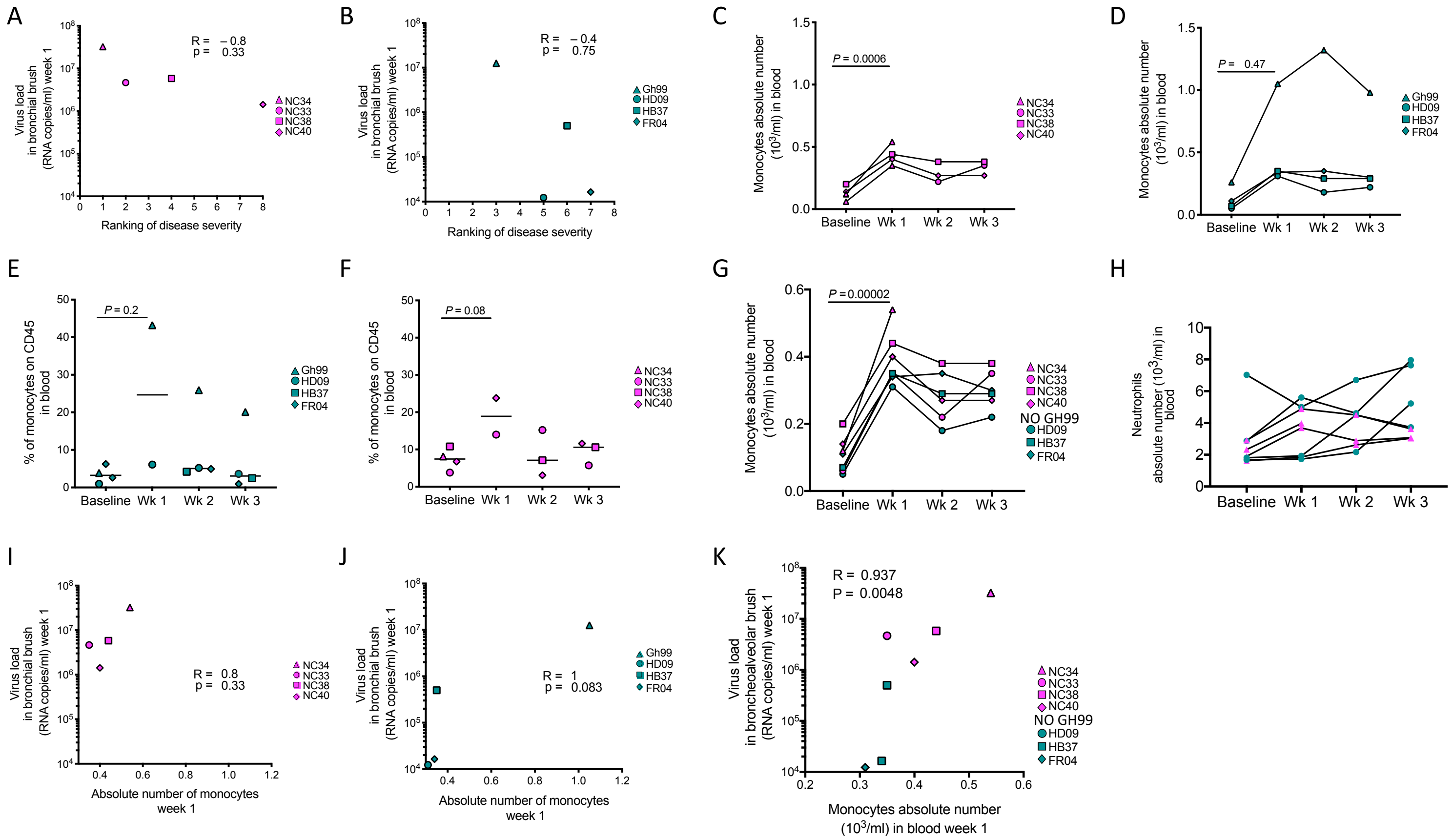


**Supplementary Figure 1. Kinetics of monocytes in rhesus macaques and African green monkeys.** Association between SARS-CoV-2 load in bronchial brushes and ranking of disease severity in AGM (**A**) and RM (**B**) (most severe case = 1, mildest case = 8). Changes in monocyte count in blood before and after infection in AGM (**C**) and RM (**D**). Proportion of total monocytes (HLA-DR+ CD16+ and/or CD14+ out of CD45+ leukocytes) in PBMCs from AGM (**E**) and RM (**F**) over time. (**G**) Absolute number of monocytes and neutrophils (**H**) in blood over time. Correlation between SARS-CoV-2 viral load in bronchial brushes and absolute count of monocytes in blood at one-week post-infection (1wk pi) in AGM (**I**) and RM (**J**). (**K**) Correlation between SARS-CoV-2 viral load in bronchial brushes and absolute count of monocytes excluding GH99 (1wk pi).

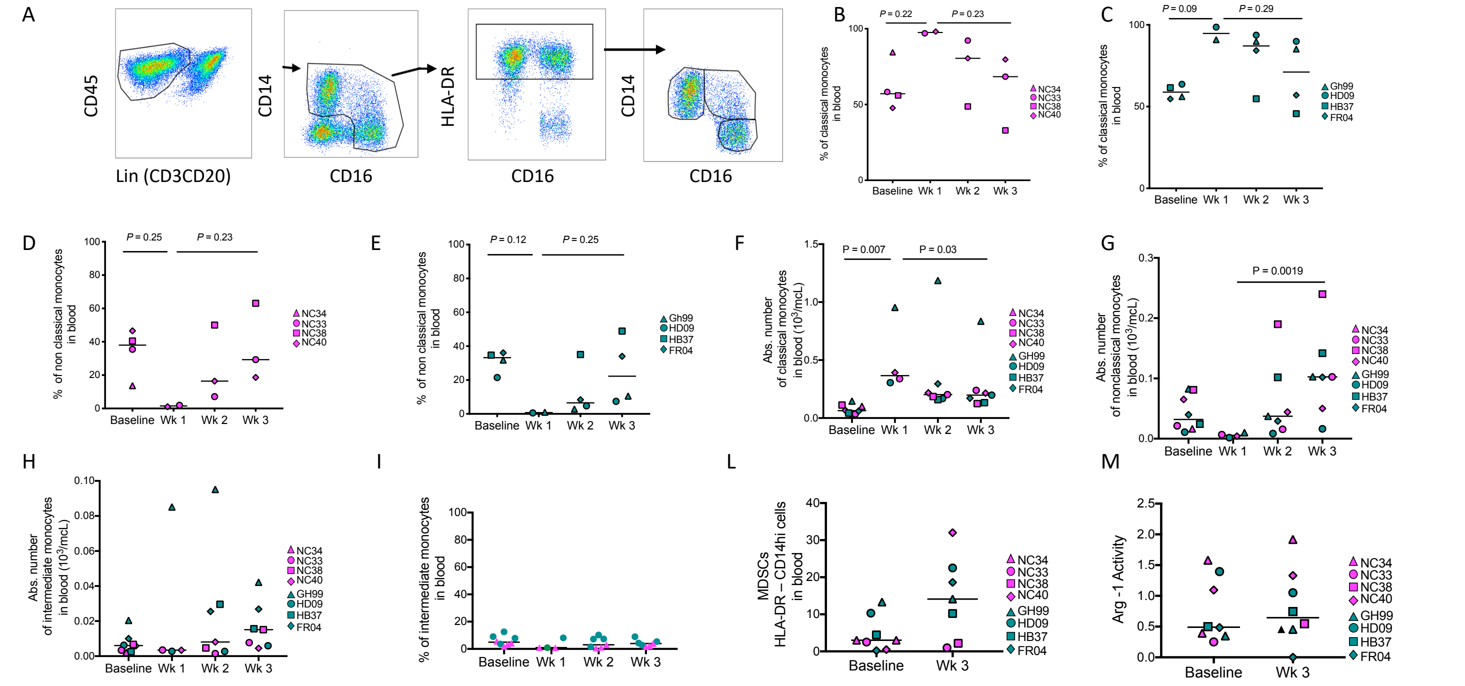


**Supplementary Figure 2. SARS-CoV-2 associated changes in monocytes subsets in blood.** (**A**) Gating strategy highlighting classification of classical (CD14hi CD16low), non-classical (CD14low CD16hi), and intermediate (CD14hi CD16hi) monocytes from blood. Percent of (**B**) classical monocytes out of total monocytes in PBMCs from AGM or RM (**C**). Percent of non-classical monocytes out of total monocytes in PBMCs in AGM (**D**) or RM (**E**). Absolute number of classical (**F**), nonclassical (**G**), and intermediate (**H**) monocytes in blood over time. **(I)** Percent of intermediate monocytes out of total monocytes in PBMCs over time. (**L**) Percent of MDSCs (HLA-DR- CD14hi) cells in blood out of live/CD45+ cells. (**M**) Arg-1 activity (unit/ml) measured at baseline or three weeks post-infection.

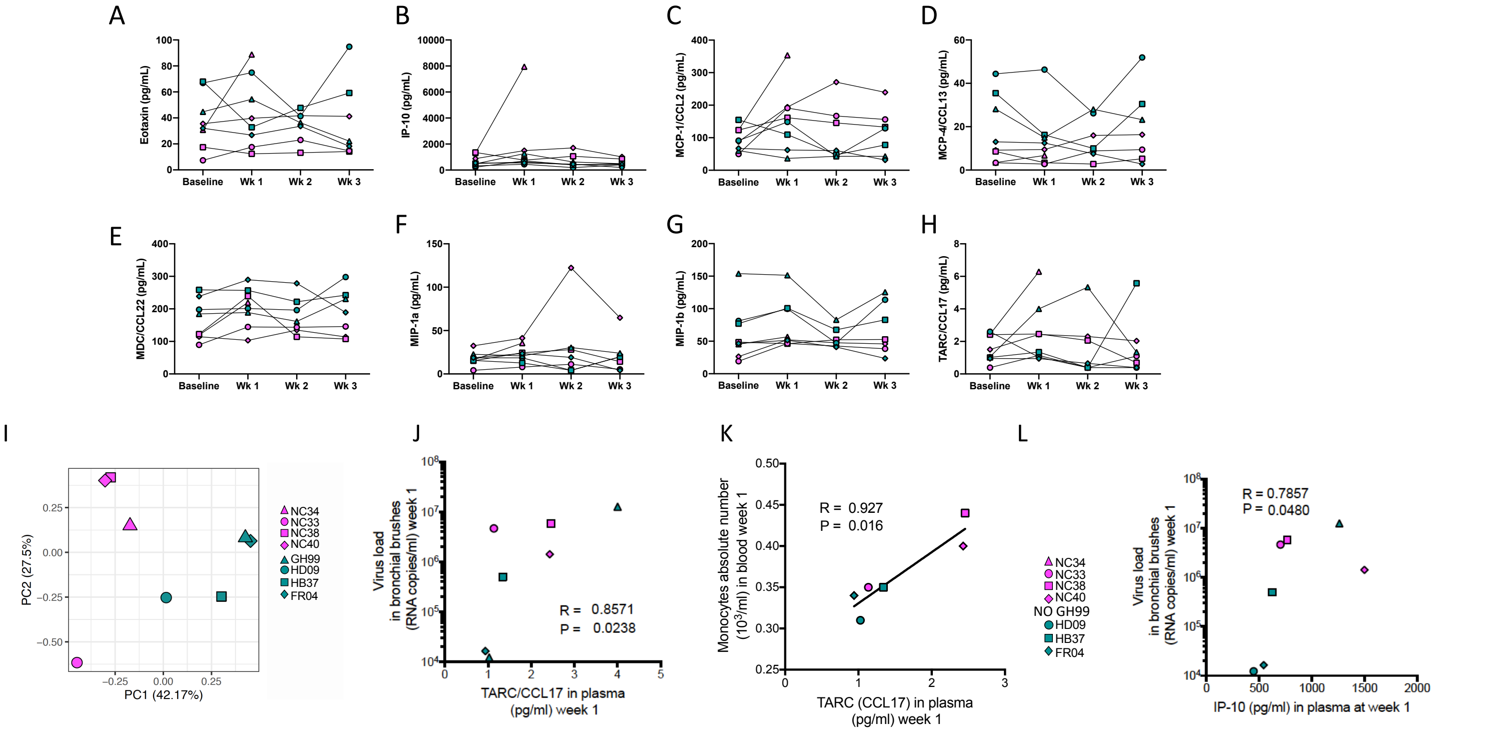


**Supplementary Figure 3**. **Chemokines promote cell migration of monocytes from the blood to the lung following infection**. Levels of chemokines in the peripheral blood over time, including (A) Eotaxin, (B) IP-10, (C) MCP-1 (CCL2), (D) MCP-4 (CCL13), (E) MDC (CCL22), (F) MIP-1a, (G), MIP-1b, and (H) TARC (CCL17). (I) Principal component analysis (PCA) plot where each point represents an individual animal and points are colored by species (AGM = magenta and RM = teal). Baseline raw data for the eight chemokines shown in A-H were used to calculate the variance between individuals that determined the position of each individual point on the plot. Principal components 1 and 2 (PC1 and PC2) explain approximately 70% of the total variance between individuals and individuals which are more closely related are clustered nearer to each other. (J) Correlation between TARC (CCL17) levels in blood and SARS-CoV-2 viral load in bronchial brushes one week post-infection. (K) Correlation between TARC (CCL17) levels in blood and the absolute number of monocytes at one week post infection. (L) Correlation between IP-10 levels in blood and SARS-CoV-2 viral load in bronchial brushes one week post-infection.

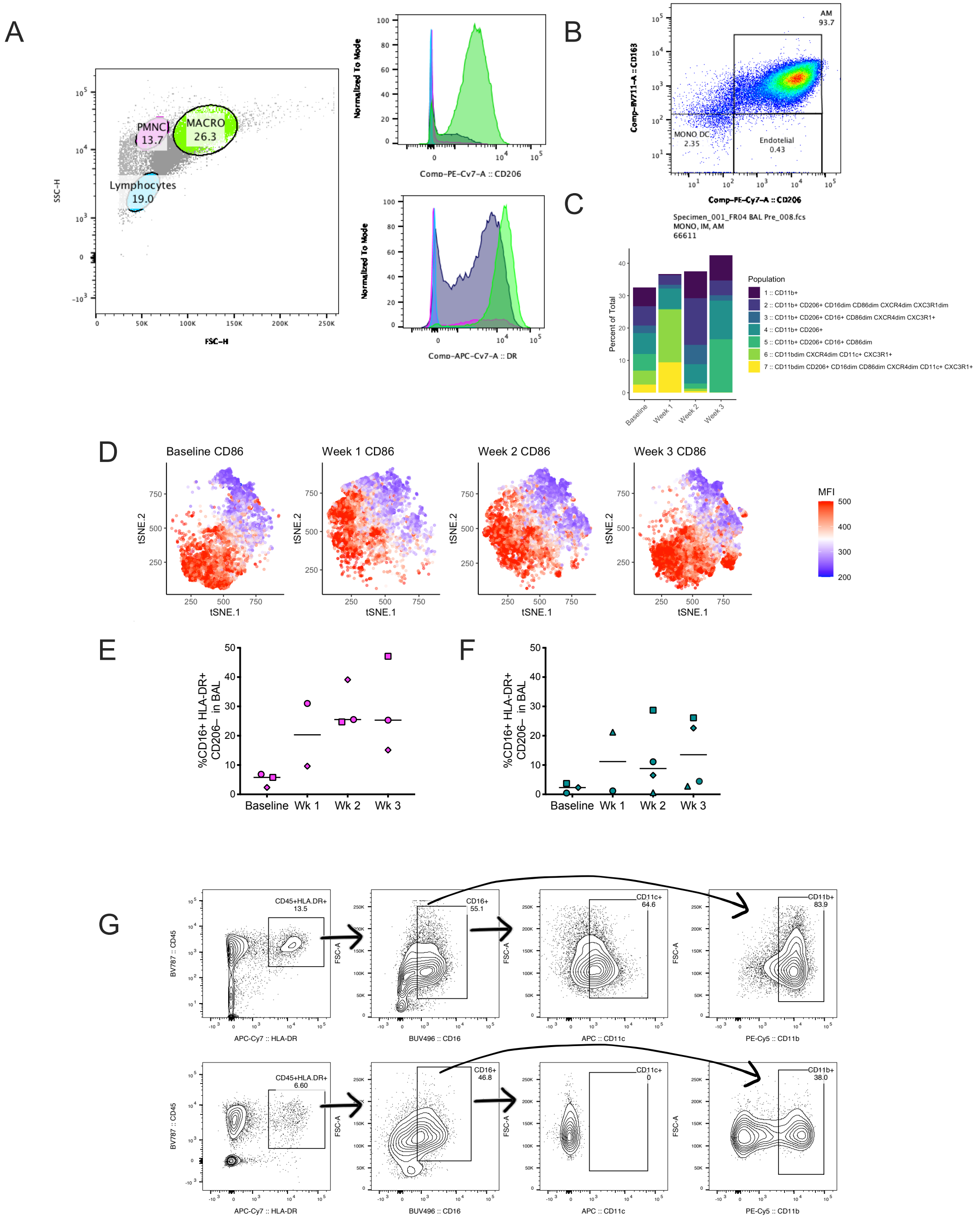


**Supplementary Figure 4. Changes in the frequency of alveolar macrophages and infiltrates in BAL during infection by flow cytometric analysis.** (**A**) Left panel. Representative light scatter (forward scatter and side scatter) analysis of lymphocytes, macrophages, and polymorphonuclear cells (PMNC) in BAL. Cell populations are located according to their relative size (forward scatter) and granularity (side scatter). Right panels. Overlaid histograms showing fluorescent intensities of CD206 and HLA-DR of lymphocytes, macrophages, and PMNC as shown on the left panels. (**B**) Representative pseudocolor dot plot displaying alveolar macrophages, endothelial cells, monocytes, and/or dendritic cells distinguished by differential expression of CD163 and CD206. (**C**) Range of phenotypes of CD11b+ and CD11c+ myeloid cells in BAL and their change in proportion out of total HLA-DR+ cells over time. (**D**) tSNE plot displaying kinetics of CD86 expression on total HLA-DR+ cells after infection. Percent of CD16+ HLA-DR+ CD206- cells in BAL over time in AGM (**E**) and RM (**F**). (**G**) Representative gating strategy showing classification of CD45+HLA-DR+CD16+ myeloid cells which express either CD11c or CD11b. Top panel showing a sample with CD11c+ cells, and bottom panel showing a sample without CD11c+ cells used to help determine optimal gate position.

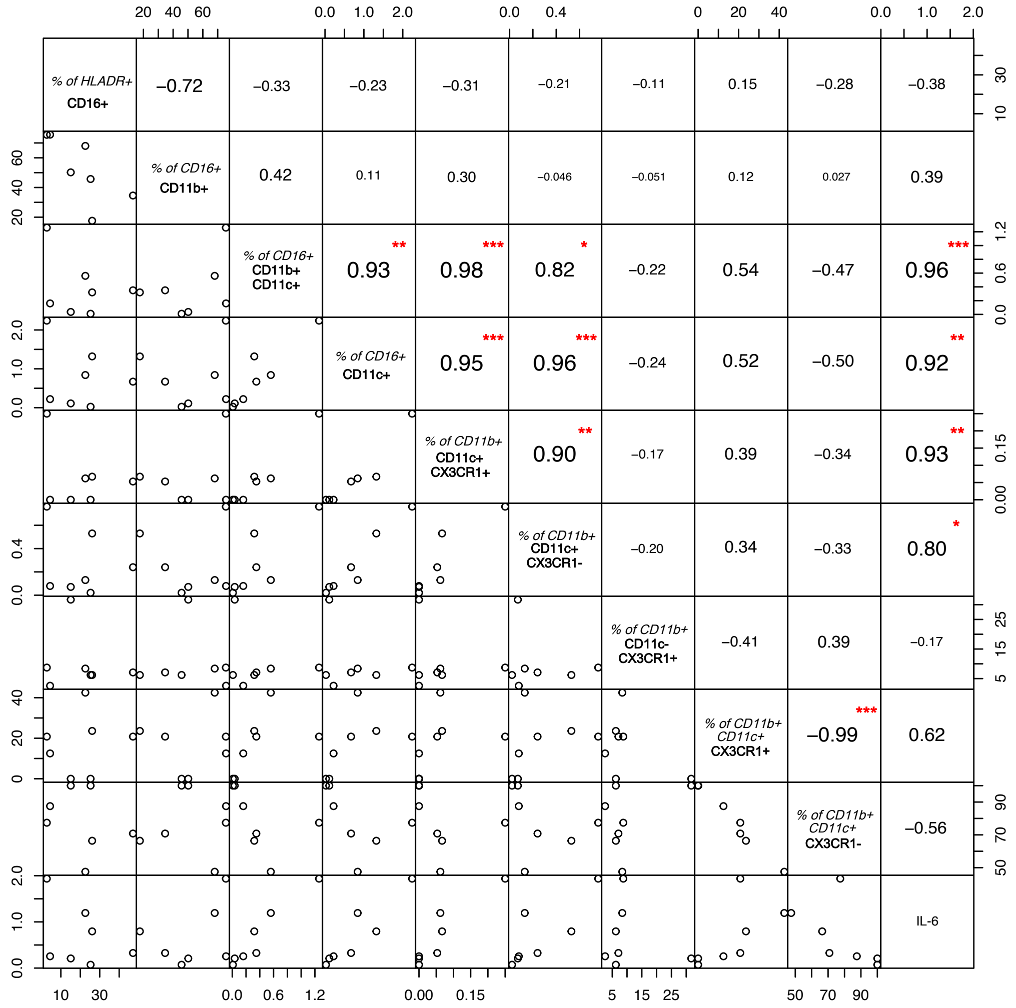


**Supplementary Figure 5**. **Association of myeloid subsets in the BAL with IL-6 plasma levels**. Correlation plot between myeloid subsets in BAL and levels of IL-6 (pg/mL) in the blood 3 weeks post infection using Pearson's method. Numbers inside boxes on the diagonal right half of the correlation plot represent the coefficient of correlation (r) and asterisks denote significance. One red asterisk corresponds with p <. 05, two red asterisks correspond with p < .01, and three red asterisks correspond with p < .001. Correlation plots in the bottom left display the actual data points.

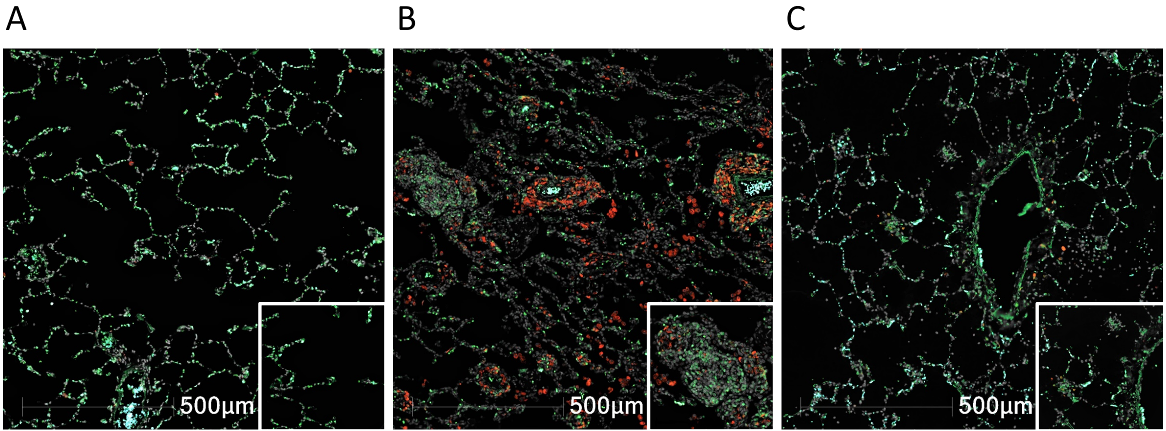


**Supplementary Figure 6. Macrophages and CD3 T cells aggregates in the lungs in infected animals.** (**A**) FR04: Isolated CD3+ cells scattered throughout the lung. (**B**) GH99: Cell aggregate with mixed population of CD3+ and CD68/CD163+ cells in isolated area of inflammation. (**C**) NC34: Rare, small aggregates of CD3+ cells (White: DAPI, CD68/CD163:red and CD3: blue

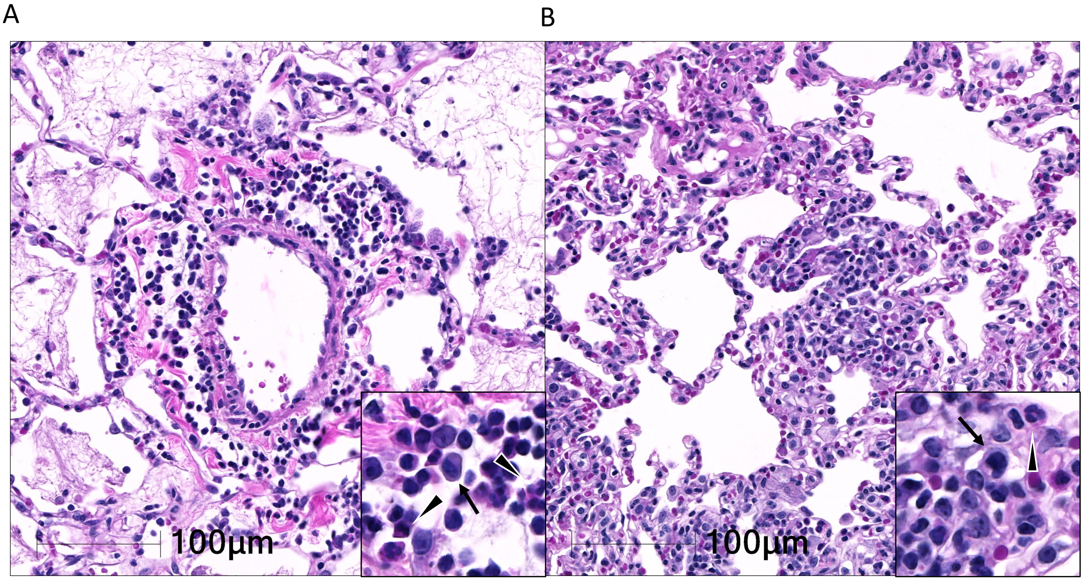


**Supplementary Figure 7. Localization of lymphocytic aggregates in SARS-CoV-2 infection**. Lymphocytes aggregates within the pulmonary interstitium. (**A**) in NC34 (necropsy day 8 post infection; rank = 1, severe) lymphocytic aggregates are perivascular and intermixed with neutrophils and histiocytes. (**B**) In NC38 (necropsy day 28, rank 5, moderate) lymphocytic aggregates are scattered within the alveolar septa and predominately mixed with histiocytes and rare neutrophils. Insets showing the admixture of macrophages (arrows) and neutrophils (arrowheads).

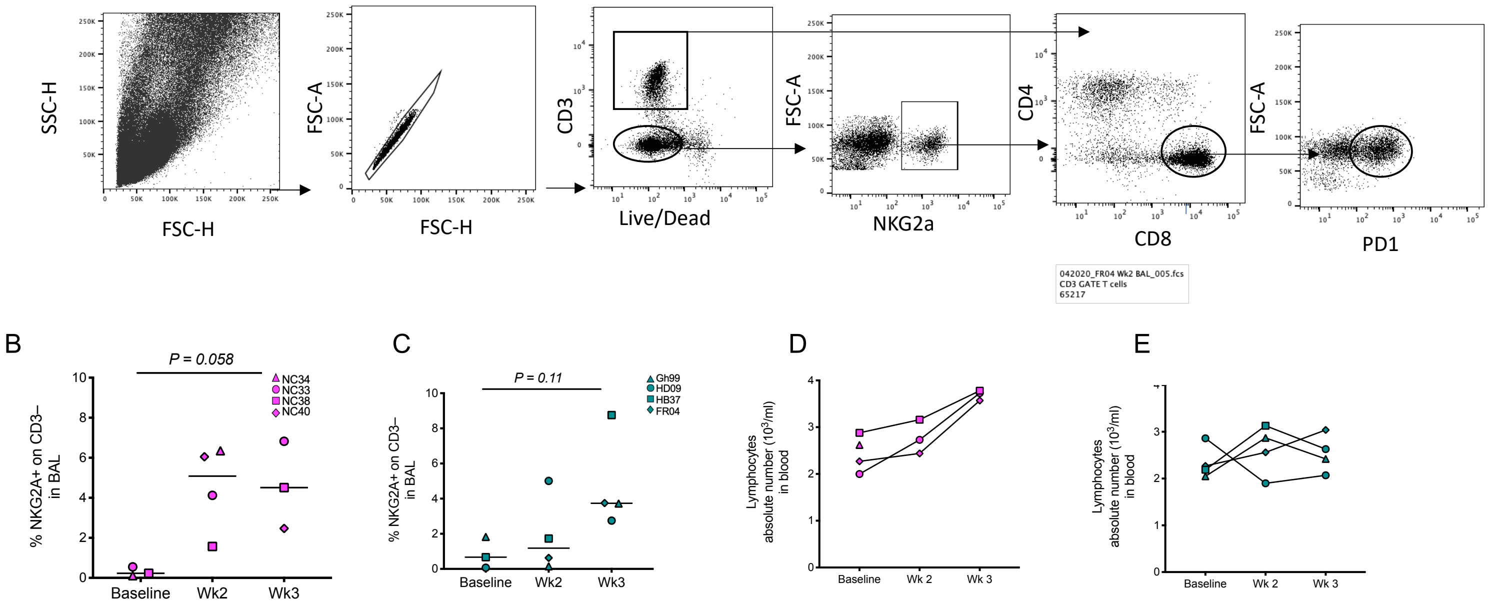


**Supplementary Figure 8. NKG2a+ cells and PD-1+ CD8 T cells.** (**A**) Gating strategy showing animal FR04 BAL at week 2. BAL was enriched of lymphocytes and NKG2a+ cells and CD8 were gated on the lymphocyte region, singled live CD3 population NKG2a+ cells were then assessed in the CD3 negative gate while CD8 T cells were gated on CD3 positive cells. The frequency of PD-1 was then assessed within the CD8 population. Percent of NKG2a+ on CD3– cells in BAL in AGM (**B**) and RM (**C**). Changes in lymphocyte count in blood before and after infection in AGM (**D**) and RM **(E**).

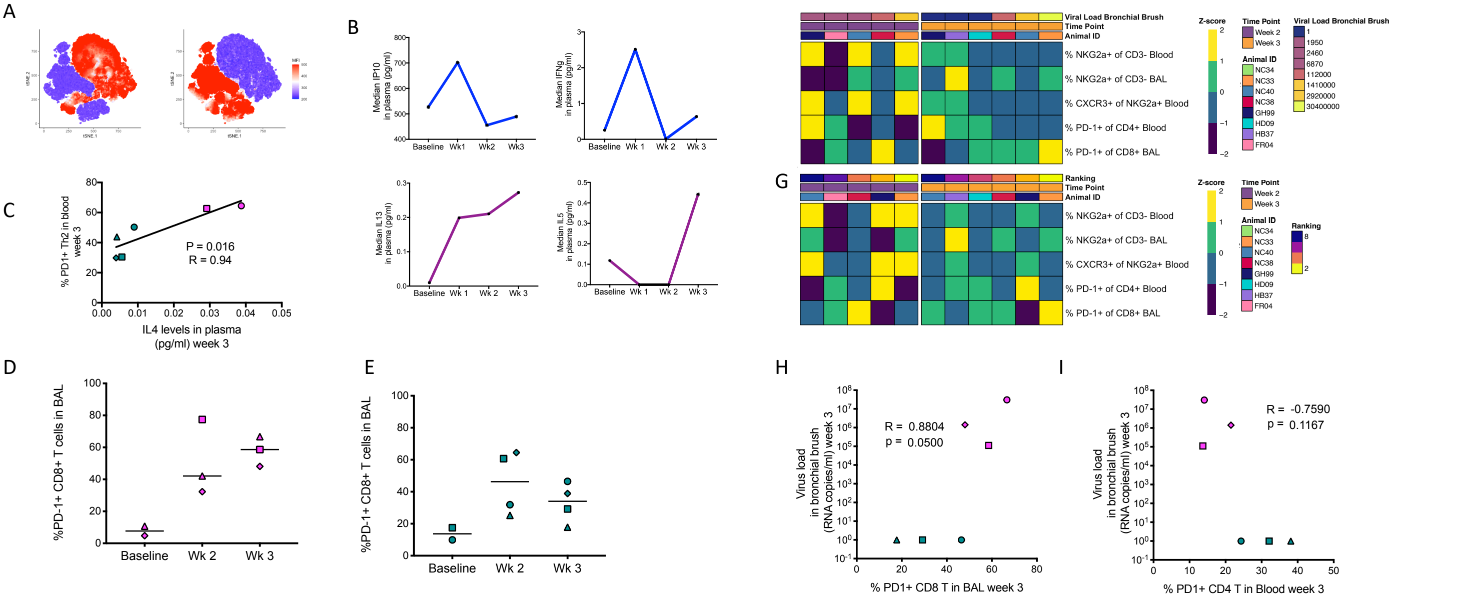


**Supplementary Figure 9. SARS-Cov2 associated changes in the quantity and the quality of T cells and NKG2a cells**(**A**) CD4 or CD8 T cells were gated from CD3 T cells from all animals over time and data were concatenated. The concatenated data were used to generate tSNE plots depicting regions where CD4 T cells (left panel) and CD8 T cells (right panel) are located. Red coloring indicates the respective location of each cell type. (**B**) Median levels of IP-10, IFN-g, IL-13, and IL-5 in plasma (pg/mL) from all eight animals over time. (**C**) Correlation between IL-4 levels in the plasma (pg/mL) and percent of PD-1+ Th2 cells in the blood at week 3 post infection. Proportion of PD-1+ CD8 T cells in BAL from AGM (D) and RM (E) after infection. (**F**) Heatmap showing the scaled frequencies of NKG2a+ and T cell subtypes in relation to viral load in bronchial brushes at weeks two (left panel) and three (right panel) post infection. The Z-score was used to determine distance from the mean and scale coloring. (**G**) Heatmap showing the scaled frequencies of NKG2a+ cells and T cell subtypes in relation to disease ranking at weeks two (left panel) and three (right panel) post infection. The Z-score was used to determine distance from the mean and scale coloring. Correlations between proportion of PD-1+ CD8 T cells (**H**) or PD-1+ CD4 T cells (**I**) and viral load in bronchial brushes three weeks post infection are shown.

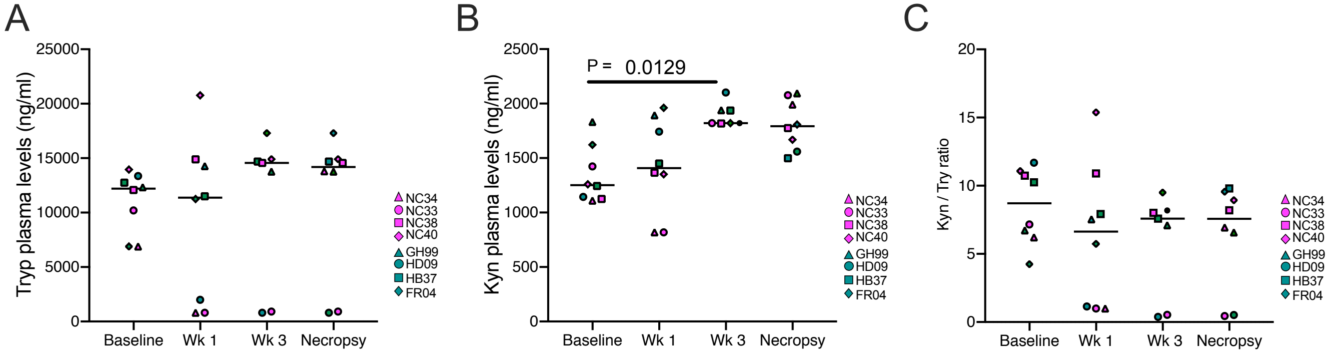


**Supplementary Figure 10. SARS-C\oV-2 associated changes in IDO activity in plasma over time.** (**A**) Levels of tryptophan (Tryp) (**B**) and Kynurenine (Kyn) (ng/ml) in plasma at week 1 and 3 post infection and at necropsy. (**C**) Kyn/Tryp ratio as a measurement of IDO activity.
