## Supplementary Tables for "Cellular events of acute, resolving or progressive COVID-19 in SARS-CoV-2 infected non-human primates"

**Supplementary Table 1**. Routes of exposure and viral inoculation

| Animal ID | Exposure (dose) | Age | Sex | Species | Base Weight (kg) |
|---|---|---|---|---|---|
| GH99 | Cumulative dose of $3.61 \times 10^6$ PFU including oral, intratracheal, intranasal, and conjunctival (both eyes) | 13.9 | Male | *Macaca mulatta*, Indian ancestry | 15.5 |
| HD09 | | 12.87 | Female | *Macaca mulatta*, Indian ancestry | 7.7 |
| NC40 | | 16 (est) | Male | *Chlorocebus aethiops* | 7.0 |
| NC33 | | 16 (est) | Female | *Chlorocebus aethiops* | 3.7 |
| HB37 | Aerosol (approx. $2.0 \times 10^3$ $TCID_{50}$) | 12.96 | Male | *Macaca mulatta*, Indian ancestry | 11.9 |
| FR04 | | 14.93 | Male | *Macaca mulatta*, Indian ancestry | 12.3 |
| NC34 | | 16 (est) | Female | *Chlorocebus aethiops* | 4.3 |
| NC38 | | 16 (est) | Male | *Chlorocebus aethiops* | 7.4 |

Est = estimated. Age of monkeys NC33, NC34, NC38, and NC40 are approximate.

**Supplementary Table 2.** Virus loads in clinical specimens

*Week 1*

| Specimen | HB37 | FR04 | NC38 | NC34 | NC33 | NC40 | GH99 | HD09 |
|---|---|---|---|---|---|---|---|---|
| *Bronchial Brush* | 4.99E+05 | 1.64E+04 | 5.81E+06 | 3.22E+07 | 4.65E+06 | 1.42E+06 | 1.26E+07 | 1.23E+04 |
| *Buccal Swab* | 3.67E+03 | 4.74E+04 | ND | 1.66E+06 | ND | 1.07E+04 | I | 2.55E+03 |
| *Nasal Swab* | 1.41E+06 | 1.52E+08 | 1.94E+04 | 6.45E+07 | 1.29E+05 | 1.05E+07 | 4.42E+04 | 1.42E+09 |
| *Pharyngeal Swab* | 5.03E+04 | 2.12E+06 | 2.47E+11 | 2.36E+09 | 7.00E+06 | 1.15E+08 | I | 6.54E+06 |
| *Rectal Swab* | 3.81E+03 | 9.90E+05 | 4.95E+03 | 4.32E+03 | 1.73E+07 | 6.22E+08 | 8.22E+03 | ND |
| *Vaginal Swab* | NT | NT | NT | 3.03E+05 | 5.16E+04 | NT | NT | ND |

*Week 2*

| Specimen | HB37 | FR04 | NC38 | NC34 | NC33 | NC40 | GH99 | HD09 |
|---|---|---|---|---|---|---|---|---|
| *Bronchial Brush* | I | 2.46E+03 | 2.16E+04 | NT | 2.92E+06 | 6.87E+03 | 1.95E+03 | I |
| *Buccal Swab* | NT | 1.71E+04 | I | NT | 6.40E+03 | 6.95E+03 | 3.78E+04 | ND |
| *Nasal Swab* | NT | 5.78E+06 | 2.07E+04 | NT | 3.27E+06 | 9.54E+04 | 1.03E+05 | 2.89E+03 |
| *Pharyngeal Swab* | 3.77E+04 | 3.11E+03 | 5.61E+04 | NT | 1.45E+03 | 9.21E+04 | 4.65E+03 | I |
| *Rectal Swab* | 2.63E+04 | 1.55E+08 | 4.69E+05 | NT | 3.93E+09 | 1.11E+05 | 2.71E+04 | ND |
| *Vaginal Swab* | NT | NT | NT | NT | 8.92E+04 | NT | NT | ND |

*Week 3*

| Specimen | HB37 | FR04 | NC38 | NC34 | NC33 | NC40 | GH99 | HD09 |
|---|---|---|---|---|---|---|---|---|
| *Bronchial Brush* | ND | I | 1.12E+05 | NT | 3.04E+07 | 1.41E+06 | ND | ND |
| *Buccal Swab* | ND | I | I | NT | ND | ND | ND | ND |
| *Nasal Swab* | ND | 7.53E+04 | 2.67E+04 | NT | 2.77E+05 | 1.42E+07 | 2.03E+04 | 4.13E+04 |
| *Pharyngeal Swab* | NT | ND | ND | NT | 1.05E+04 | 8.95E+04 | I | 4.28E+04 |
| *Rectal Swab* | ND | 1.31E+04 | 3.78E+06 | NT | 1.45E+04 | 3.45E+04 | ND | ND |
| *Vaginal Swab* | NT | NT | NT | NT | I | NT | NT | ND |

ND = Not Determined, NT = Not Tested due to lack of sample availability, I = Inconclusive.

**Supplementary Table 3.** Temperature and oxygen saturation.

| | *Baseline* | | *Week 1* | | *Week 2* | | *Week 3* | |
|---|---|---|---|---|---|---|---|---|
| **Animal ID** | **Temperature** | **$SpO_2$** | **Temperature** | **$SpO_2$** | **Temperature** | **$SpO_2$** | **Temperature** | **$SpO_2$** |
| FR04 | 100.4 | 98 | 100.9 | 93 | 101.1 | 95 | 102.3 | 94 |
| GH99 | 100.4 | 99 | 101.5 | 97 | 100.3 | 92 | 100.8 | 93 |
| HB37 | 100.9 | 91 | 102.6 | 94 | 102.2 | 95 | 103.5 | 96 |
| HD09 | 101.1 | 98 | 100.7 | 95 | 101.3 | 93 | 101.3 | 94 |
| NC33 | 100.3 | - | 99.5 | 96 | 100.1 | 96 | 99.4 | 92 |
| NC34 | 99.9 | 96 | 100.9 | 90 | - | - | - | - |
| NC38 | 99.9 | 94 | 102.2 | 96 | 101.2 | 95 | 100.8 | 91 |
| NC40 | 102.1 | 94 | 101.4 | 97 | 101.4 | 98 | 102.1 | 94 |

Temperature measured in °F.

**Supplementary Table 4**. Comparison of disease outcomes by route of challenge or sex.

*Aerosol vs. Multi-route (p-values adjusted)*

| Disease Factor | Baseline | Week 1 | Week 2 | Week 3 | Necropsy |
|---|---|---|---|---|---|
| *$SpO_2$* | 0.5743 | 0.2387 | 0.9998 | 0.9986 | - |
| *Temperature* | 0.7245 | 0.5392 | 0.7532 | 0.244 | - |
| *Viral Load* | - | 0.9025 | 0.9996 | 0.6114 | - |
| *Ranking* | - | - | - | | >.9999 |

*Female vs. Male (p-values adjusted)*

| Disease Factor | Baseline | Week 1 | Week 2 | Week 3 | Necropsy |
|---|---|---|---|---|---|
| *$SpO_2$* | 0.8436 | 0.7922 | 0.9985 | 0.9969 | - |
| *Temperature* | 0.9809 | 0.1554 | 0.914 | 0.1556 | - |
| *Viral Load* | - | 0.514 | 0.987 | 0.0504 | - |
| *Ranking* | - | - | - | | 0.1012 |

P-values were adjusted for multiple comparisons. Data for rankings were based only on the necropsy time point.

**Supplementary Table 5**. Correlations between viral loads from different specimens and disease ranking.

| | *Week 1* | | | *Week 2* | | | *Week 3* | | |
|---|---|---|---|---|---|---|---|---|---|
| **Specimen** | **R** | **P-value** | **P-value (adj)** | **R** | **P-value** | **P-value (adj)** | **R** | **P-value** | **P-value (adj)** |
| *Bronchial brush* | -0.71 | 0.058 | 0.754 | -0.3 | 0.683 | 1 | -0.5 | 1 | 1 |
| *Buccal swab* | -0.1 | 0.95 | 1 | 0.2 | 0.917 | 1 | - | - | - |
| *Nasal swab* | 0.31 | 0.462 | 1 | -0.14 | 0.803 | 1 | 0.43 | 0.419 | 1 |
| *Pharyngeal swab* | -0.46 | 0.302 | 1 | 0.6 | 0.242 | 1 | 1 | 0.333 | 1 |
| *Rectal swab* | 0.36 | 0.444 | 1 | -0.26 | 0.658 | 1 | - | - | - |
| *Vaginal swab* | - | - | - | - | - | - | - | - | - |

R = correlation coefficient. P-values were adjusted for multiple comparisons. Missing statistical calculations are due to lack of sample availability or inconclusive results.

**Supplementary Table 6.** Correlations between viral loads from bronchial brushes and the frequency and absolute number of monocyte subtypes.

| Cell type | Week 1 | | | Week 2 | | | Week 3 | | |
|---|---|---|---|---|---|---|---|---|---|
| | R | P-value | P-value (adj) | R | P-value | P-value (adj) | R | P-value | P-value (adj) |
| % Classical monocytes | -0.9867 | 0.0133 | 0.0798 | 0.3823 | 0.5254 | 1 | 0.0409 | 0.9387 | 1 |
| % Nonclassical monocytes | 0.1767 | 0.8233 | 1 | -0.2534 | 0.6809 | 1 | -0.0162 | 0.9757 | 1 |
| % Intermediate monocytes | 0.9430 | 0.057 | 0.342 | -0.5848 | 0.3004 | 1 | -0.4343 | 0.3895 | 1 |
| Absolute number classical monocytes | 0.9393 | 0.0607 | 0.3642 | -0.2921 | 0.6335 | 1 | -0.1015 | 0.8483 | 1 |
| Absolute number nonclassical monocytes | 0.9627 | 0.0373 | 0.2238 | -0.3379 | 0.5781 | 1 | -0.0548 | 0.9179 | 1 |
| Absolute number intermediate monocytes | 0.9407 | 0.0593 | 0.3558 | -0.3795 | 0.5287 | 1 | -0.2807 | 0.5901 | 1 |

R = correlation coefficient.

**Supplementary Table 7.** Levels of plasma chemokines.

| Animal ID | Time Point | Eotaxin | IP-10 | MCP-1 | MCP-4 | MDC | MIP-1a | MIP-1b | TARC |
|---|---|---|---|---|---|---|---|---|---|
| *FR04* | *Baseline* | 32.01 | 547.00 | 67.13 | 13.04 | 238.50 | 18.65 | 46.56 | 0.95 |
| | *Week 1* | 26.75 | 542.34 | 62.31 | 12.45 | 289.45 | 23.62 | 51.40 | 0.94 |
| | *Week 2* | 33.64 | 455.34 | 60.43 | 7.55 | 278.59 | 19.06 | 41.03 | 0.65 |
| | *Week 3* | 18.17 | 196.14 | 31.44 | 2.79 | 189.15 | 4.23 | 23.66 | 0.39 |
| | *Necropsy* | 22.69 | 364.97 | 62.48 | 7.50 | 254.21 | 15.74 | 40.65 | 0.42 |
| *GH99* | *Baseline* | 44.71 | 482.21 | 60.30 | 28.10 | 184.46 | 22.66 | 153.92 | 1.05 |
| | *Week 1* | 54.40 | 1260.46 | 36.66 | 14.96 | 188.69 | 20.86 | 151.48 | 4.00 |
| | *Week 2* | 36.33 | 620.06 | 42.93 | 28.03 | 161.50 | 30.38 | 82.88 | 5.33 |
| | *Week 3* | 22.14 | 491.42 | 42.77 | 23.22 | 230.56 | 23.94 | 125.41 | 1.34 |
| | *Necropsy* | 33.72 | 452.24 | 57.13 | 20.75 | 326.46 | 19.89 | 143.02 | 1.33 |
| *HB37* | *Baseline* | 67.97 | 506.65 | 154.69 | 35.46 | 258.36 | 15.53 | 77.16 | 1.01 |
| | *Week 1* | 32.83 | 622.46 | 109.66 | 16.21 | 256.82 | 12.72 | 100.79 | 1.34 |
| | *Week 2* | 47.69 | 408.33 | 47.34 | 9.98 | 221.84 | 4.23 | 67.51 | 0.39 |
| | *Week 3* | 59.22 | 489.13 | 77.68 | 30.49 | 242.08 | 19.83 | 82.99 | 5.57 |
| | *Necropsy* | 45.32 | 630.00 | 104.97 | 16.66 | 518.60 | 13.95 | 105.09 | 1.34 |
| *HD09* | *Baseline* | 66.89 | 324.83 | 91.16 | 44.36 | 197.82 | 18.52 | 81.26 | 2.60 |
| | *Week 1* | 74.87 | 447.65 | 148.55 | 46.33 | 201.15 | 18.59 | 99.71 | 1.03 |
| | *Week 2* | 41.40 | 187.00 | 42.77 | 26.16 | 196.32 | 4.23 | 47.31 | 0.39 |
| | *Week 3* | 94.82 | 407.99 | 128.95 | 51.95 | 297.78 | 19.66 | 113.81 | 0.39 |
| | *Necropsy* | 66.19 | 541.69 | 93.97 | 34.73 | 371.65 | 17.53 | 104.06 | 1.39 |
| *NC33* | *Baseline* | 7.39 | 221.91 | 49.81 | 3.35 | 89.52 | 4.23 | 19.19 | 0.39 |
| | *Week 1* | 17.54 | 701.70 | 191.19 | 2.79 | 144.63 | 7.93 | 46.54 | 1.14 |
| | *Week 2* | 23.03 | 404.76 | 166.82 | 8.84 | 143.51 | 11.22 | 42.59 | 0.39 |
| | *Week 3* | 14.65 | 436.50 | 156.17 | 9.42 | 145.98 | 5.26 | 38.46 | 1.10 |
| | *Necropsy* | 13.78 | 185.31 | 295.92 | 2.79 | 71.28 | 4.23 | 56.30 | 0.39 |
| *NC34* | *Baseline* | 30.78 | 1303.10 | 123.93 | 3.45 | 119.71 | 16.19 | 45.48 | 2.44 |
| | *Week 1* | 88.83 | 7943.91 | 353.42 | 6.75 | 219.50 | 35.56 | 56.68 | 6.29 |
| | *Necropsy* | 116.82 | 78897.78 | 1606.75 | 6.00 | 300.12 | 29.53 | 232.02 | 6.23 |
| *NC38* | *Baseline* | 17.49 | 1357.14 | 123.61 | 8.60 | 122.65 | 15.16 | 48.62 | 2.42 |
| | *Week 1* | 12.33 | 766.14 | 161.39 | 3.35 | 239.25 | 24.47 | 46.26 | 2.46 |
| | *Week 2* | 13.22 | 1065.47 | 145.35 | 2.79 | 114.31 | 28.07 | 52.18 | 2.05 |
| | *Week 3* | 14.13 | 845.43 | 132.72 | 5.21 | 107.54 | 14.18 | 52.82 | 0.69 |
| | *Necropsy* | 31.00 | 794.69 | 184.65 | 15.35 | 156.91 | 28.71 | 33.34 | 0.79 |
| *NC40* | *Baseline* | 35.58 | 875.23 | 87.90 | 9.15 | 114.59 | 32.46 | 26.43 | 1.50 |
| | *Week 1* | 39.70 | 1498.56 | 194.17 | 9.45 | 103.31 | 41.35 | 52.20 | 2.43 |
| | *Week 2* | 41.68 | 1703.38 | 270.96 | 15.97 | 134.91 | 122.34 | 47.64 | 2.30 |
| | *Week 3* | 41.24 | 1014.66 | 239.48 | 16.33 | 113.86 | 64.80 | 45.91 | 2.03 |
| | *Necropsy* | 37.49 | 1611.59 | 185.71 | 15.12 | 149.16 | 60.97 | 49.18 | 3.08 |

Values are shown in pg/mL

**Supplementary Table 8.** $Log_2$ fold change of chemokines relative to baseline.

| Animal ID | Time Point | Eotaxin | IP-10 | MCP-1 | MCP-4 | MDC | MIP-1a | MIP-1b | TARC |
|---|---|---|---|---|---|---|---|---|---|
| *FR04* | *Week 1* | -0.26 | -0.01 | -0.11 | -0.07 | 0.28 | 0.34 | 0.14 | -0.02 |
| | *Week 2* | 0.07 | -0.26 | -0.15 | -0.79 | 0.22 | 0.03 | -0.18 | -0.55 |
| | *Week 3* | -0.82 | -1.48 | -1.09 | -2.23 | -0.33 | -2.14 | -0.98 | -1.29 |
| | *Necropsy* | -0.50 | -0.58 | -0.10 | -0.80 | 0.09 | -0.24 | -0.20 | -1.17 |
| *GH99* | *Week 1* | 0.28 | 1.39 | -0.72 | -0.91 | 0.03 | -0.12 | -0.02 | 1.92 |
| | *Week 2* | -0.30 | 0.36 | -0.49 | 0.00 | -0.19 | 0.42 | -0.89 | 2.34 |
| | *Week 3* | -1.01 | 0.03 | -0.50 | -0.28 | 0.32 | 0.08 | -0.30 | 0.35 |
| | *Necropsy* | -0.41 | -0.09 | -0.08 | -0.44 | 0.82 | -0.19 | -0.11 | 0.34 |
| *HB37* | *Week 1* | -1.05 | 0.30 | -0.50 | -1.13 | -0.01 | -0.29 | 0.39 | 0.40 |
| | *Week 2* | -0.51 | -0.31 | -1.71 | -1.83 | -0.22 | -1.88 | -0.19 | -1.38 |
| | *Week 3* | -0.20 | -0.05 | -0.99 | -0.22 | -0.09 | 0.35 | 0.11 | 2.46 |
| | *Necropsy* | -0.58 | 0.31 | -0.56 | -1.09 | 1.01 | -0.15 | 0.45 | 0.40 |
| *HD09* | *Week 1* | 0.16 | 0.46 | 0.70 | 0.06 | 0.02 | 0.01 | 0.30 | -1.34 |
| | *Week 2* | -0.69 | -0.80 | -1.09 | -0.76 | -0.01 | -2.13 | -0.78 | -2.74 |
| | *Week 3* | 0.50 | 0.33 | 0.50 | 0.23 | 0.59 | 0.09 | 0.49 | -2.74 |
| | *Necropsy* | -0.02 | 0.74 | 0.04 | -0.35 | 0.91 | -0.08 | 0.36 | -0.90 |
| *NC33* | *Week 1* | 1.25 | 1.66 | 1.94 | -0.27 | 0.69 | 0.91 | 1.28 | 1.55 |
| | *Week 2* | 1.64 | 0.87 | 1.74 | 1.40 | 0.68 | 1.41 | 1.15 | 0.00 |
| | *Week 3* | 0.99 | 0.98 | 1.65 | 1.49 | 0.71 | 0.31 | 1.00 | 1.49 |
| | *Necropsy* | 0.90 | -0.26 | 2.57 | -0.27 | -0.33 | 0.00 | 1.55 | 0.00 |
| *NC34* | *Week 1* | 1.53 | 2.61 | 1.51 | 0.97 | 0.87 | 1.13 | 0.32 | 1.37 |
| | *Necropsy* | 1.92 | 5.92 | 3.70 | 0.80 | 1.33 | 0.87 | 2.35 | 1.35 |
| *NC38* | *Week 1* | -0.50 | -0.82 | 0.38 | -1.36 | 0.96 | 0.69 | -0.07 | 0.02 |
| | *Week 2* | -0.40 | -0.35 | 0.23 | -1.63 | -0.10 | 0.89 | 0.10 | -0.24 |
| | *Week 3* | -0.31 | -0.68 | 0.10 | -0.72 | -0.19 | -0.10 | 0.12 | -1.81 |
| | *Necropsy* | 0.83 | -0.77 | 0.58 | 0.84 | 0.36 | 0.92 | -0.54 | -1.62 |
| *NC40* | *Week 1* | 0.16 | 0.78 | 1.14 | 0.05 | -0.15 | 0.35 | 0.98 | 0.69 |
| | *Week 2* | 0.23 | 0.96 | 1.62 | 0.80 | 0.24 | 1.91 | 0.85 | 0.62 |
| | *Week 3* | 0.21 | 0.21 | 1.45 | 0.84 | -0.01 | 1.00 | 0.80 | 0.44 |
| | *Necropsy* | 0.08 | 0.88 | 1.08 | 0.72 | 0.38 | 0.91 | 0.90 | 1.04 |

**Supplementary Table 9.** Rotation table derived from PCA plots.

| | PC1 | PC2 | PC3 | PC4 | PC5 | PC6 | PC7 | PC8 |
|---|---|---|---|---|---|---|---|---|
| Eotaxin | 0.51349854 | 0.14431892 | 0.04547296 | 0.10908504 | -0.2471479 | 0.22538446 | -0.6843669 | -0.3482608 |
| IP-10 | -0.2522775 | 0.55999126 | -0.0765231 | 0.02243312 | 0.41068394 | -0.1618914 | 0.05487606 | -0.6469353 |
| MCP-1 | 0.11727127 | 0.51945467 | 0.4952681 | 0.25731776 | 0.20002956 | 0.44221279 | 0.14981145 | 0.38328048 |
| MCP-4 | 0.51986265 | -0.0023534 | 0.07196805 | -0.2206943 | -0.2665758 | 0.11724746 | 0.67825244 | -0.3619612 |
| MDC | 0.45844363 | -0.0508976 | 0.21350047 | 0.3735466 | 0.24485256 | -0.7283914 | 0.0253733 | 0.10473167 |
| MIP-1a | 0.15406887 | 0.2517465 | -0.7797945 | 0.47174206 | -0.1062296 | 0.09007799 | 0.16325863 | 0.19030037 |
| MIP-1b | 0.39295357 | -0.0473738 | -0.2926367 | -0.510066 | 0.65448152 | 0.14743768 | -0.1017211 | 0.19263585 |
| TARC | 0.01858282 | 0.57230412 | -0.0480451 | -0.5004337 | -0.3994979 | -0.39152 | -0.0922837 | 0.31301387 |

All elements used for the Principal Component (PC) Analysis are shown. PC = PCA Coefficients for each case (row = each chemokine) on each factor (column) that can be plotted. The square of each PC defines the amount of variation contributed by each variable within the factor.

**Supplementary Table 10**. Antibodies used for flow cytometry and immunohistochemistry.

*Flow cytometry antibodies*

| Marker | Fluorochrome | Clone | Ref. | Company | Known Reactivity |
|---|---|---|---|---|---|
| PD-1 | APC | EH12.2H7 | 329908 | Biolegend | RM & AGM |
| CD56 | Alexa 700 | B159 | 557919 | BD | RM _ AGM not reported |
| CD183 (CXCR3) | APC/Cyanine7 | G025H7 | 353722 | Biolegend | RM & AGM |
| CD196 (CCR6 ) | BV605 | G034E3 | 353420 | Biolegend | RM & AGM |
| CD4 | BV797 | L200 | 563914 | BD | RM & AGM |
| CD3 | PE | SP34-2 | 552127 | BD | RM & AGM |
| CD3 | BV650 | SP34-2 | 563918 | BD | RM & AGM |
| CD20 | BV650 | 2H7 | 563780 | BD | RM & AGM |
| CD95 | PE/Cyanine5 | DX2 | 559773 | BD | RM & AGM |
| CD8a | Texas Red | RPA-T8 | MHCD0817 | Life technology | RM & AGM |
| CD159a (NKG2a) | PE/Cyanine7 | Z199 | B10246 | Beck Coulter | RM & AGM |
| CD16 | BV711 | 3G8 | 563127 | BD | RM & AGM |
| CD16 | BUV496 | 3G8 | 564653 | BD | RM & AGM |
| CD86 | FITC | FUN-1 | 557343 | BD | RM _ AGM not reported |
| CD11c | APC | 3.9 | 17-0116-42 | eBioscence | RM & AGM |
| HLA-DR | APC/Cyanine7 | L243 | 307618 | Biolegend | RM & AGM |
| CD14 | V450 | M5E2 | 561390 | BD | RM & AGM |
| CD163 | BV711 | GHI/61 | 563889 | BD | RM _ AGM not reported |
| CD184 (CXCR4) | PECF594 | 12G5 | 562389 | BD | RM & AGM |
| CD45 | BV786 | D058-1283 | 563861 | BD | RM & AGM |
| CX3CR1 | PE | K0124E1 | 355704 | Biolegend | RM _ AGM not reported |
| CD11b | PE/Cyanine5 | ICRF44 | 301308 | Biolegend | RM & AGM |
| CD206 | PE/Cyanine7 | 19.2 | 25-2069-42 | BD | RM & AGM |

*Immunohistochemistry antibodies*

| Primary Antibody | Species | Company | Ref. | Dilution | Secondary Antibody |
|---|---|---|---|---|---|
| CD206 | Goat | R&D | AF2535 | 1:50 | Donkey anti-goat 647 (far red) |
| CD11b | Rabbit | Abcam | Ab133357 | 1:20 | Donkey anti-rabbit 488 (green) |
| CD16 | Mouse IgG2a | Novocastra | NCL-CD16 | 1:40 | Permanent Red |
| DAPI | Dye - N/A | Invitrogen | D1306 | 1:20,000 | N/A |
| MPO | Rabbit | Dako | A0398 | 1:6000 | Permanent Red |
| CD3 | Rabbit | Dako | A0452 | 1:50 | Goat anti-rabbit 488 (green) |
| CD68 | Mouse IgG1 | Dako | M0814 | 1:20 | Goat anti-IgG1 568 (red) |
| CD163 | Mouse IgG1 | Leica | NCL-L-CD163 | 1:50 | Goat anti-IgG1 568 (red) |
